# Horizontal transfer of chromosomal DNA mediated by an integrative and conjugative element generates frequent localized recombination in *Novosphingobium aromaticivorans*

**DOI:** 10.64898/2026.03.24.713927

**Authors:** Marco N. Allemann, Leah H. Hochanadel, Delyana P. Vasileva, Joshua K. Michener

## Abstract

Horizontal gene transfer is an important evolutionary process by which DNA is exchanged between cells that are physically co-located but not direct evolutionary descendants. Horizontal transfer of highly divergent DNA is relatively easy to detect and can produce major phenotypic changes, exemplified by acquisition of antibiotic resistance determinants. However, transfer of high-identity DNA, for example between strains of the same species, is likely to be more frequent, harder to detect, and highly impactful in aggregate. In this work, we demonstrate that recombination between soil isolates of the alphaproteobacterium *Novosphingobium aromaticivorans* can exchange chromosomal DNA, leading to multiple unselected recombination events spanning approximately 10% of the chromosome. Chromosomal recombination was directional, more efficient near an integrative and conjugative element (ICE), and required a relaxase found in the ICE. Recombination could not be observed into strains from closely related *Novosphingobium* species. In combination, these results suggest that ICE-mediated recombination can efficiently recombine DNA within *N. aromaticivorans*, increasing the adaptive potential of the species while also enforcing species boundaries through preferential intraspecific recombination.

**Importance:** Horizontal gene transfer is a key process in bacterial evolution. Mechanisms for transfer of mobile genetic elements are well-characterized, but less is known about how chromosomal DNA is recombined. In this work, we demonstrate that integrative and conjugative elements can efficiently recombine chromosomal DNA between strains of *Novosphingobium aromaticivorans* but not between different *Novosphingobium* species. We conclude that ICE-mediated chromosomal recombination can be an important adaptive mechanism within a species, due to its ability to recombine nearby chromosomal alleles, but also serves to delineate species-specific gene pools as a result of its limited phylogenetic range.

## Introduction

Horizontal gene transfer (HGT) is a major evolutionary mechanism in microbes (1). Many examples of HGT are generated by transfer of distinct mobile genetic elements (MGEs) such as plasmids, and these MGEs frequently carry genes encoding highly-impactful genetic factors such as antibiotic resistance markers or pathogenicity determinants (2). Transfer of genomic DNA, particularly between closely-related strains, is also thought to occur frequently, but is harder to detect through comparative genomics (3, 4). Intraspecific recombination is likely an important mechanism in bacterial speciation and the maintenance of species-specific gene pools (5–7).

Horizontal transfer of genetic material between bacteria has been demonstrated to occur via diverse mechanisms including conjugation, transduction, and transformation, and these mechanisms can transfer both chromosomal and extrachromosomal DNA (8, 9). For example, the F plasmid of *Escherichia coli* can integrate into the chromosome and mobilize adjacent DNA through a process known as ‘high frequency recombination’ or Hfr (10). Other examples of conjugative mobilization of genomic DNA include distributive conjugative transfer in *Mycobacteria*, wherein numerous, unlinked fragments of chromosomal DNA can be transferred in a single conjugative event (11). Additional bacterial genera such as *Mycoplasma (12), Yersinia (13), Helicobacter (14)*, and *Vibrio* (15) have also been shown to transfer genomic DNA via conjugation between relatively closely related strains. However, more general mechanisms are likely responsible for the majority of intraspecific DNA exchange.

Integrative and Conjugative Elements (ICEs) are a group of MGEs that replicate while site-specifically integrated into the host chromosome but then excise, circularize, and transfer into neighboring cells by conjugation (16, 17). While conjugative plasmids have been well-studied, ICEs are thought to be at least as abundant in bacterial genomes (18). ICEs primarily transfer themselves or associated ‘integrative and mobilizable elements’ and can disseminate associated genetic cargo. However, in a process similar to plasmid-mediated Hfr, ICEs can initiate conjugation prior to excision, leading to transfer of adjacent chromosomal DNA (19). Given the abundance of ICEs and their ability to transfer chromosomal DNA, ICE-mediated Hfr has the potential to serve as a general mechanism for chromosomal exchange.

Relatively little is known about recombination after Hfr, whether mediated by an ICE or a chromosomally-integrated plasmid. Traditional mating experiments have demonstrated transfer of multiple loci and inferred genetic distances based on the likelihood of simultaneous transfer (20). However, chromosome-wide nucleotide-level analysis of the resulting transconjugants has only recently been described. Reported examples of chromosomal recombination through conjugative transfer between strains of *E. coli* (21) or *Bacteriodes fragilis* (22) have described recombination primarily of a single contiguous segment of DNA. A more recent approach in *E. coli* using engineered Hfr measured variable rates of recombination, with progeny inheriting between one and sixteen DNA fragments from the donor (23). In addition to its evolutionary role in the environment, the ability to recombine large segments of chromosomal DNA makes Hfr-mediated genetic exchange a useful tool for strain construction and analysis (24–26). Understanding the likely genomic consequences of Hfr-mediated recombination can guide the design and application of these approaches.

Many studies on bacterial HGT have been conducted with model strains such as *E. coli* or pathogens and other host-associated microbes. However, the rhizosphere can support dense microbial populations and therefore are important sites for HGT (27). Sphingomonads are Gram-negative alphaproteobacteria found abundantly in the rhizosphere that have been extensively studied for their abilities to degrade a wide range of anthropogenic and naturally-occurring aromatic compounds (28). Among these sphingomonads, *Novosphingobium aromaticivorans* F199 (hereafter ‘F199’) has recently emerged as a promising model organism for discovery of novel lignin degradation pathways and lignin valorization to bioproducts (29–36). The genome of F199 has been sequenced and consists of the main chromosome (3.3Mb) and two large plasmids designated pNL1 (184 Kb) and pNL2 (500 kb). The plasmid pNL1 was previously sequenced and was shown to contain regions related to plasmid replication, conjugal mobilization, and catabolism of aromatic compounds (37). F199 was isolated from a subsurface borehole at the US Department of Energy Savannah River Site (38). Two other *N. aromaticivorans* strains were isolated contemporaneously from a separate borehole at the same site (39) but have not been characterized to the same degree as F199.

In this work, we characterized genomic recombination between strains of *N. aromaticivorans*. We demonstrated that recombination is directional, with one strain consistently contributing a minority of the progeny genome, and is mediated by an ICE present in the minor parent. Recombination was efficient between strains of *N. aromaticivorans* but was not detected in closely-related *Novosphingobium* strains. Recombination was localized to approximately 10% of the genome, but chimeric progeny often contained multiple independent recombination events within this window. We also observed ICE-independent recombination between mutants of *N. aromaticivorans* F199 and demonstrated that these recombination events were not mediated by plasmid-dependent Hfr. In combination, these results demonstrate that ICE-mediated Hfr in *N. aromaticivorans* leads to efficient intraspecific recombination, thereby enhancing evolution within the species while contributing to interspecific genetic segregation.

## Materials and Methods

### Bacterial strains and growth conditions

*Novosphingobium* strains were routinely grown at 30 °C in R2A medium. Minimal defined media for *Novosphingobium* was medium 457 (DSMZ) with 0.2% glucose as a sole carbon source. *Escherichia coli* strains were grown in Luria-Bertani (LB) medium. For solid media, agar was included at 15 g/L. The antibiotics kanamycin (50 µg/ml), streptomycin (100 µg/ml), chloramphenicol (15 µg/ml) were used as required. The amino acid supplements histidine (100 µM) and leucine (300 µM) were added to defined media as needed. When necessary, diaminopimelic acid (DAP) was used at 60 µg/ml.

### Mutagenesis in *Novosphingobium aromaticivorans*

Mutants of *N. aromaticivorans* strains were generated by allele exchange mutagenesis using the plasmid pAK405 as described previously (29, 40). Resistance cassettes for kanamycin or chloramphenicol were cloned between homologous regions (500-700 bp) upstream and downstream of the gene to be replaced. The sequence for the kanamycin cassette was derived from the pAK405 backbone (40). The chloramphenicol resistance cassette was designed by placing the chloramphenicol resistance gene from pRL271 (41) under the control of the artificial sphingomonad psyn2 promoter described previously (40). All pAK405 knockout constructs were synthesized by Genscript or assembled using HiFi Assembly (New England Biolabs, Ipswich, MA) following manufacturer recommendations.

Derivatives of pAK405 were mobilized into *N. aromaticivorans* via biparental conjugation using *E*.*coli* WM6026, a DAP auxotroph, as described previously (29). Selection for exconjugants was performed on R2A + kanamycin or R2A + chloramphenicol agar at 30 °C. Selections were performed in the absence of DAP to counterselect against the WM6026 donor strain. Exconjugants were clonally purified by restreaking to the same selection media. Exconjugants were grown overnight in R2A broth without antibiotics and plated to R2A with streptomycin supplemented with 2.5 g/L NaCl to counterselect against the integrated plasmid. Streptomycin-resistant colonies were screened by patching for appropriate amino acid auxotrophies on 457 media with 2 g/L glucose. Gene replacements were further verified with colony PCR and whole genome resequencing.

### Novosphingobium conjugations

*Novosphingobium* strains were grown overnight in R2A broth at 30°C with the appropriate antibiotic selection(s). The next day, cells were pelleted and washed with R2A broth to remove residual antibiotics and equivalent cell numbers, as calculated by OD600, of washed cells from each strain were mixed together, pelleted, and concentrated into a minimal volume which was then spotted onto the surface of a dry R2A agar plate. Once the conjugation mixture dried, it was incubated at 30 °C for 24 h. After incubation, cells were scraped off the surface of the agar plate and resuspended in R2A media lacking carbon. For conjugations that required double antibiotic selection, resuspended cells were plated to R2A containing the appropriate combination of antibiotics. For conjugations that required auxotrophic counterselection against one of the strains, the resuspended cells were washed a minimum of three times, by centrifugation at 10,000 x g for 1 min at room temperature and resuspension in carbon-source-free 457 media, before plating to selective media to remove nutrient carryover.

| Strain | Description | Source |
| --- | --- | --- |
| *Escherichia coli* strains |  |  |
| WM6026 | *lacIq*, *rrnB3*, Δ*lacZ*4787, *hsdR514*, Δ*araBAD*567, Δ*rhaBAD*568, *rph-1*, *attλ::pAE12*(Δ*oriR6K-cat::Frt*5), Δ*endA::Frt*, *uidA*(Δ*MluI*)::*pir*, *attHK::pJK1006D*(*oriR6K-cat::Frt*5; *trfA::Frt*) *dap* | (42) |
| DH5α | *fhuA2 lac(del)U169 phoA glnV44 Φ80' lacZ(del)M15 gyrA96 recA1 relA1 endA1 thi-1 hsdR17* | Lab stock |
| *Novosphingobium* strains |  |  |
| F199 | Wild-type strain, DSM12444 | DSMZ |
| B0522 | *Novosphingobium* sp. B0522 | (39) |
| B0695 | *Novosphingobium* sp. B0695 | (39) |
| F199C | F199, cured of pNL1 | This study |
| JMN150 | F199, Δ*hisB::kan* | (25) |
| JMN127 | F199, Δ*leuB::cat* | (25) |
| JMN154 | F199, Δ*hisB::kan*, Δ*leuB::cat* | (25) |
| JMN146 | F199C, Δ*hisB::kan* | This study |
| JMN149 | F199C, Δ*leuB::cat* | This study |
| JMN155 | F199C, Δ*hisB::kan*, *leuB::cat* | This study |
| JMN71 | B0522 Δ*hisB::kan* | This study |
| JMN228 | B0695 Δ*hisB::kan* | This study |
| *Escherichia coli* strains |  |  |
| JMN218 | B0695 $\Delta leuB::cat$ | This study |
| JMN219 | B0695 $\Delta hisB::kan \Delta leuB::cat$ | This study |
| JMN229 | B0695 $\Delta rel::cat$ | This study |
| Plasmids |  |  |
| pAK405 | Allele exchange plasmid for sphingomonads, Kan<sup>R</sup>, *rpsL* (Sm<sup>S</sup>) | (40) |
| pJM613 | pAK405, F199 $\Delta hisB::kan$ | (25) |
| pJM614 | pAK405, F199 $\Delta leuB::cat$ | (25) |
| pJM | pAK405, B0695 $\Delta rel::cat$ | This work |

## Results and Discussion

### Characterization of intraspecific recombination in *N. aromaticivorans*

We previously generated recombinant strains of *N. aromaticivorans* using a modified protoplast fusion protocol (43). In the protoplast fusion process, the outer membrane and peptidoglycan are removed through chemical and enzymatic treatment to generate cells surrounded by only a single membrane (44). Protoplasts can then be chemically fused, providing an opportunity for genome-wide recombination (43). Crossing F199 Δ*hisB*::*kan* with F199 Δ*leuB*::*cat*, followed by selection for kan^R^ cm^R^ progeny, readily yielded a F199 Δ*hisB*::*kan* Δ*leuB*::*cat* strain, as confirmed by genome resequencing. Crossing F199 Δ*hisB*::*kan* Δ*leuB*::*cat* with B0695, followed by selection for kan^R^ leu^+^ progeny, yielded chimeric progeny strains. Genome resequencing demonstrated that these chimeric progeny inherited the majority of their genomes from F199, including the Δ*hisB*::*kan* marker, but recombined different segments of DNA from B0695, including the wild-type *leuB* allele (25). Unselected DNA that was introduced from B0695 was primarily observed near the *leuB* locus, in a region spanning approximately 400 kbp.

While optimizing the recombination process, we occasionally observed recombinant progeny in our control crosses, which used parental strains that had not been treated with lysozyme. While these progeny were generated at lower rates than those from lysozyme-treated parents, resequencing identified similar patterns of localized recombination near the selection marker. Since progeny from untreated parents could not be the result of protoplast fusion, we investigated other mechanisms that could yield large-scale genome recombination.

To further characterize the lysozyme-independent recombination mechanism, we repeated the same B0695 x F199 Δ*hisB*::*kan* Δ*leuB*::*cat* cross, but without lysozyme treatment of the parents. We split the resulting cell mixture and selected for progeny with either kan^R^ leu^+^ or cm^R^ his^+^ phenotypes, since these would indicate recombination between the parental strains (Figure S1). This approach typically generated at least 10-fold more kan^R^ leu^+^ progeny than cm^R^ his^+^ progeny, consistent with different rates of recombination at the *hisB* and *leuB* loci. Similarly, recombinant kan^R^ leu^+^ progeny typically appeared within three days, while cm^R^ his^+^ colonies took more than a week to appear. Inclusion of DNAse did not affect the formation of recombinant progeny, suggesting that recombination was not due to natural competence.

We next sequenced twelve progeny for each of the phenotypes, kan^R^ leu^+^ or cm^R^ his^+^. Eleven of the twelve kan^R^ leu^+^ progeny were unique recombinants, with patterns of recombination similar to those observed previously. Only five of the twelve cm^R^ his^+^ progeny were unique; the genomes of the remaining seven were duplicates of one of these five. Regardless of the selection, F199 contributed the majority of the genomic DNA, with smaller fragments recombined from B0695 (Figure 1). The recombined fragments were located near the selection marker, either *leuB* or *hisB*.

**Figure 1:**
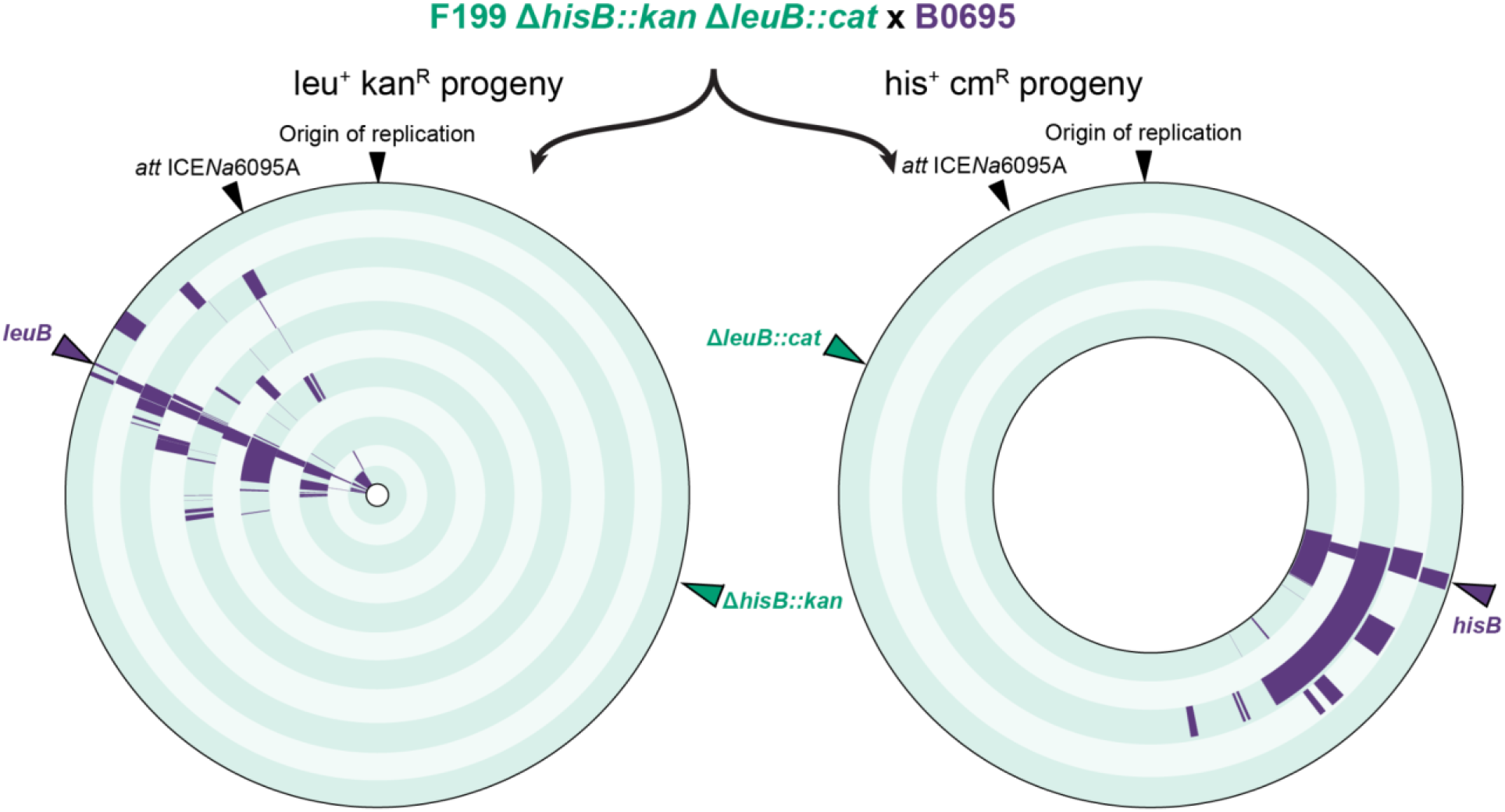
Genomes of chimeric transconjugants resulting from the F199 Δ*hisB*::*kan* Δ*leuB*::*cat* x B0695 cross. Each concentric circle represents the genome of a unique progeny strain. Segments highlighted in purple are derived from B0695. The remaining DNA is ambiguous or derived from F199. Black arrows identify the origin of replication and the integration site for ICE*Na*6095A. Green and purple arrows highlight the selection markers derived from F199 or B0695, respectively.

To test whether these results were strain-specific, we generated additional parental strains, B0695 Δ*leuB*::*cat* and B0522 Δ*hisB::kan*. Crossing B0695 *leuB::cat* with B0522 *hisB::kan* yielded kan^R^ cm^R^ progeny but not his^+^ leu^+^ progeny. Resequencing the genomes of ten kan^R^ cm^R^ progeny demonstrated that B0522 contributed the majority of the genomic DNA, including Δ*hisB::kan*, with the DNA from B0695 located near the Δ*leuB::cat* marker (Figure S2). Overall, results with B0522 were similar to those using F199, though quantitatively less efficient: B0695 contributed a minority of the progeny genomes, and the *leuB* locus was transferred more efficiently than *hisB*.

To evaluate the potential phylogenetic breadth of this mechanism for chromosomal recombination, we constructed a B0695 Δ*hisB::kan* Δ*leuB::cat* mutant and crossed it with three closely-related *Novosphingobium* species: *Novosphingobium stygium* DSM12445, *Novosphingobium capsulatum* DSM30196, and *Novosphingobium subterraneum* DSM12447 (79-83% average nucleotide identity, Figure S3). None of these crosses yielded kan^R^ leu^+^ or cm^R^ his^+^ progeny.

### Analysis of chromosomal recombination patterns

Individual homologous recombination events can be estimated based on genome resequencing. However, the precision of this approach depends on the frequency of variable nucleotide positions that differentiate the donor and recipient DNA. Consistent with our previous approach (43), we measured the minimum length for each recombined segment of DNA, from the first contiguous B0695-derived variant to the last; the full length of recombined DNA could be substantially longer (Figure S4B). Additionally, we analyzed each genome as the result solely of recombination from B0695 into F199. Some of the patterns of recombination could be explained equally parsimoniously by sequential recombination of B0695 and F199 DNA, perhaps in a merodiploid intermediate(Figure S4C). B0695 differs from F199 at 24,747 nucleotide positions, producing a marker roughly every 144 bp.

Progeny resulting from crossing B0695 x F199 Δ*hisB*::*kan* Δ*leuB*::*cat* contained 1-14 B0695-derived DNA per strain (mean 5.07, median 5; Figure 2A). The distribution of fragment lengths was bimodal, with one population of small (<10 bp) fragments representing ∼18% of the total recombination events and a second population with a median length of 13.3 kbp representing the remaining 82% (Figure 2B). We note that the population of short recombination events could be an artifact of the low frequency of nucleotide markers. However, the maximum possible length of these fragments, typically 300 bp, is still substantially lower than the average length of the longer population. The total fraction of the F199 genome replaced by B0695-derived DNA varied from 0.8% to 13.7% (mean 4.0%, median 2.2%; Figure 2C). No significant differences were observed between his^+^ cat^R^ and leu^+^ kan^R^ progeny.

**Figure 2:**
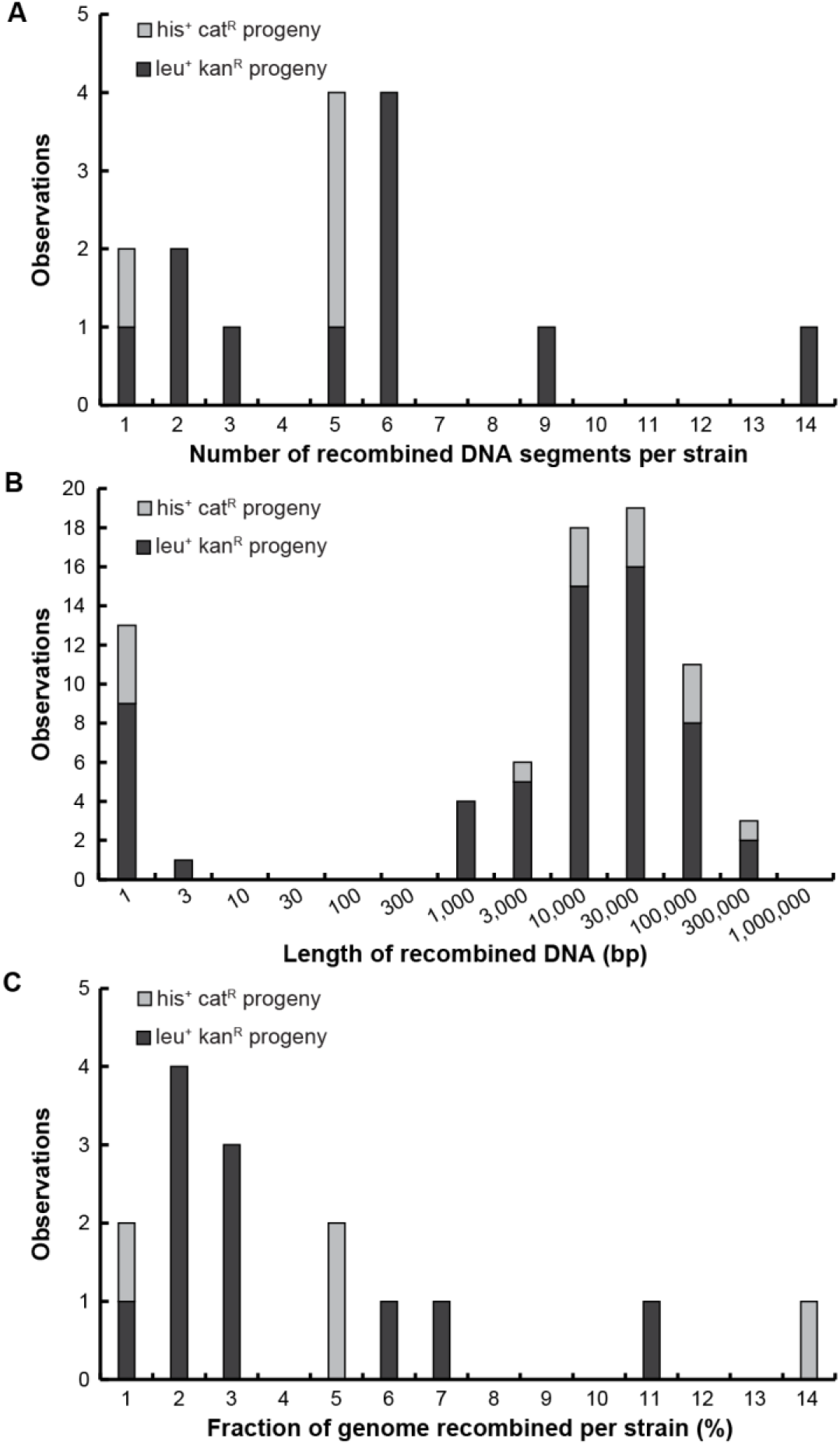
Characterization of DNA recombination during ICE-mediated Hfr. (A) The number of recombined fragments was calculated for each recombinant strain, varying from 1-14 fragments per strain with a median of 5. (B) The minimum length of each fragment was calculated as shown in Figure S4B. (C) The total length of B0695-derived fragments was summed for each strain and reported as a fraction of the genome length.

Compared to previous reports using engineered Hfr in *E. coli* (23) or protoplast fusion in *B. subtilis (25, 43)*, the numbers of recombined fragments in *N. aromaticivorans* were similar to engineered Hfr, with a mean of approximately 5 fragments per strain, but both were lower than protoplast fusion, which generated a mean of approximately 20 fragments per strain. All three methods recombined a broad range of fragment sizes. Additionally, the percentage genome recombined from a single cross was similar, approximately 2-3%. Compared to protoplast fusion, the other two methods generated higher local recombination rates across a smaller portion of the genome.

### Elucidation of recombination mechanisms

We first sought to explain recombination between F199 Δ*hisB*::*kan* and F199 Δ*leuB*::*cat*, which we used to generate the F199 Δ*hisB*::*kan* Δ*leuB*::*cat* double mutant. All three *N. aromaticivorans* strains contain homologs of pNL1 (37), a self-mobilizable plasmid that can transfer to related sphingomonads (45). We hypothesized that transient chromosomal integration of this plasmid could also mediate Hfr conjugation (46). To test this hypothesis, we used a strain of F199, known as ‘F199C’, that we generated previously during serial propagation with coumaric acid as a sole growth substrate. Resequencing F199C demonstrated that it spontaneously lost pNL1, similar to our previous reports of large deletions in pNL1 during serial propagation with related lignin-derived aromatic compounds (31, 47). Other than the loss of pNL1, no other significant mutations were observed in F199C relative to F199. Therefore, we constructed F199C Δ*hisB*::*kan* and F199C Δ*leuB*::*cat* strains and compared the efficiency of recombination between otherwise equivalent strains, F199C Δ*hisB*::*kan* x F199C Δ*leuB*::*cat* versus F199 Δ*hisB*::*kan* x F199 Δ*leuB*::*cat*. Both crosses yielded progeny at equivalent rates (Figure 3), and analysis of the F199C-derived progeny confirmed both the presence of the both resistance markers as well as the lack of pNL1. Therefore, we conclude that the mechanism for recombination between F199 mutants is not pNL1-mediated Hfr. Further analysis will be needed to identify this recombination process.

**Figure 3:**
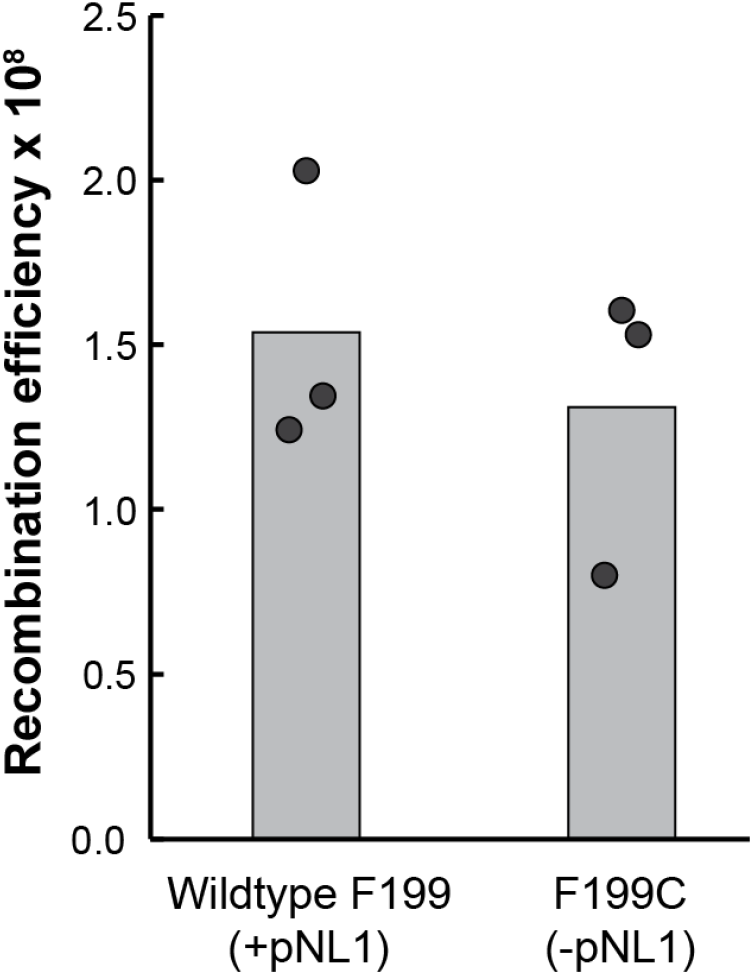
Chromosomal recombination is not mediated by the conjugative plasmid pNL1. Wildtype F199 Δ*hisB*::*kan* was crossed with F199 Δ*leuB*::*cat*, both of which contain pNL1, and compared to F199C Δ*hisB*::*kan* crossed with F199C Δ*leuB*::*cat*, both of which lack pNL1. Both crosses generated recombinant kan^R^ cm^R^ progeny at similar rates.

Therefore, we next examined potential mechanisms for directional recombination from B0695 into F199 or B0522. Comparing the genomes of B0695, B0522, and F199 identified a putative integrative and conjugative element (ICE) present in B0695 but absent from the other two strains (48). This ICE, hereafter ‘ICE*Ns*0695A’, is integrated into a leucine tRNA gene located approximately 400 kbp away from the *leuB* locus (Figure 1). Similar to previous reports in *Vibrio cholerae* (19) and *Bacteroides fragilis (22)*, we hypothesized that ICE*Ns*0695A could be responsible for efficiently mobilizing adjacent chromosomal DNA from B0695 into B0522 or F199. This hypothesis was consistent with the observed patterns of genomic recombination, since recombination of B0695 DNA was primarily observed in the genomic region between ICE*Ns*0695A and the *leuB* locus. It is unclear whether the distal *hisB* locus could be transferred through a similar ICE-dependent mechanism, but the physical separation on the chromosome could account for lower recombination efficiency. Increased rates of recombination after lysozyme treatment, as initially observed, could result from physiological stress triggering ICE-mediated conjugation.

To test this hypothesis, we constructed a strain of B0695 in which the putative relaxase in ICE*Ns*0695A was replaced with the *cat* marker. We then crossed F199 Δ*hisB*::*kan* Δ*leuB*::*cat* with either wild-type B0695 or B0695 Δ*rel*::*cat* and selected for kan^R^ leu^+^ progeny. The cross with B0695 Δ*rel*::*cat* yielded no kan^R^ leu^+^ progeny, while the cross using wild-type B0695 produced 660 kan^R^ leu^+^ cfu/mL (Figure 4). Based on these results, we concluded that recombination of DNA adjacent to the *leuB* locus from B0695 into F199 was mediated by ICE*Ns*0695A through a mechanism similar to high frequency recombination (Hfr).

**Figure 4:**
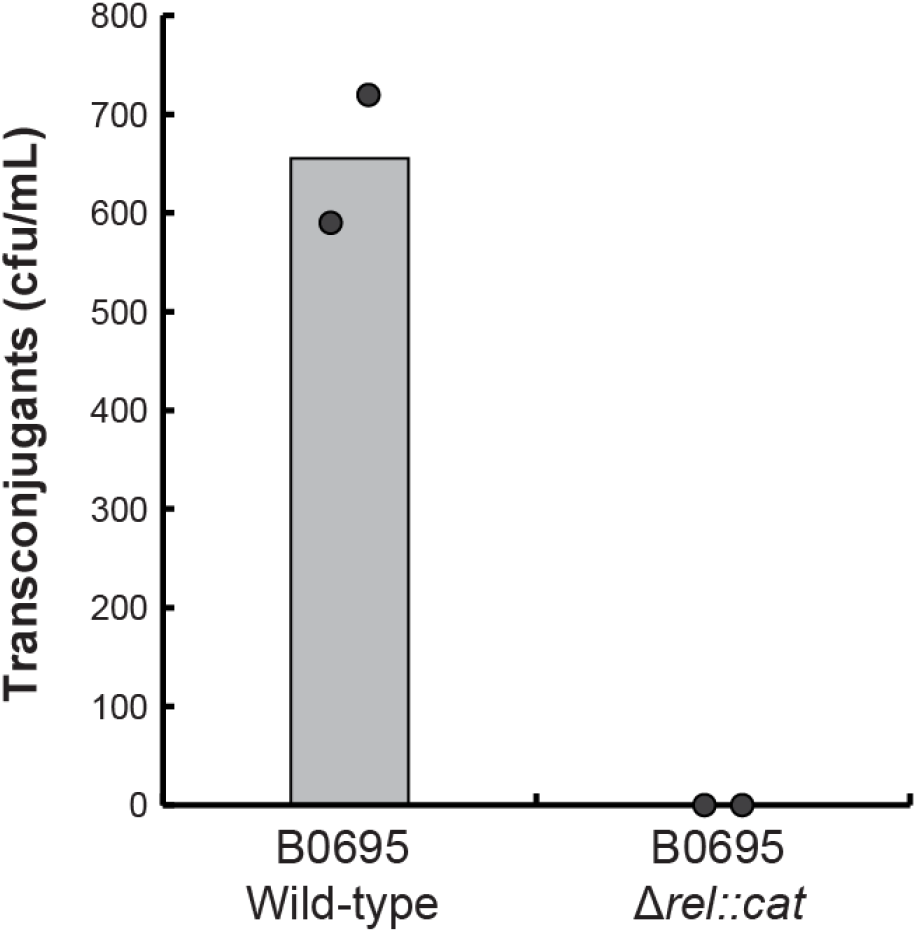
Generation of recombinant progeny requires the ICE*Ns*6095A relaxase. F199 Δ*hisB*::*kan* Δ*leuB*::*cat* was crossed with either wild-type or mutant B0695, followed by selection for leu^+^ kan^R^ progeny that had recombined the wild-type *leuB* locus from B0695. No transconjugants were observed when the relaxase mutant was used as the donor.

## Conclusion

In this work, we demonstrated that an ICE present in *N. aromaticivorans* B0695 could efficiently transfer and recombine chromosomal DNA into two other *N. aromaticivorans* strains but not into more-distantly-related *Novosphingobium* species. Multiple fragments of DNA could be transferred in a single event, spanning a broad range of lengths. Enhanced intraspecific recombination is a key process in the adaptation and maintenance of bacterial species, and our results highlight ICE-mediated HGT as a plausible mechanism for intraspecific recombination of diverse DNA sequences. This process could also be engineered to construct recombinant bacterial strains for purposes such as genetic mapping or biotechnology.

## Acknowledgements

This manuscript has been authored by UT-Battelle, LLC under Contract No. DE-AC05-00OR22725 with the U.S. Department of Energy (DOE). This work was primarily supported by the U.S. DOE, Office of Science, Office of Biological and Environmental Research, though an Early Career Award to JKM. Analysis of recombinant genomes was supported by the Center for Bioenergy Innovation, U.S. Department of Energy, Office of Science, Biological and Environmental Research Program under Award Number ERKP886 (D.P.V and J.K.M).

